# Cost-effective mapping of genetic interactions in mammalian cells

**DOI:** 10.1101/2021.04.29.441342

**Authors:** Arshad H. Khan, Desmond J. Smith

## Abstract

Comprehensive maps of genetic interactions in mammalian cells are daunting to construct because of the large number of potential interactions, ~ 2 × 10^8^ for protein coding genes. We previously used co-inheritance of distant genes from published radiation hybrid (RH) datasets to identify genetic interactions. However, it was necessary to combine six legacy datasets from four species to obtain adequate statistical power. Mapping resolution was also limited by the low density PCR genotyping. Here, we employ shallow sequencing of nascent human RH clones as an economical approach to constructing interaction maps. In this initial study, 15 clones were analyzed, enabling construction of a network with 225 genes and 2359 interactions (FDR < 0.05). Despite its small size, the network showed significant overlap with the previous RH network and with a protein-protein interaction network. Consumables were ≲ $50 per clone, showing that affordable, high quality genetic interaction maps are feasible in mammalian cells.

## Introduction

Intelligent intervention in normal and diseased mammalian cells requires a comprehensive map of their biological networks. Protein-protein interactions (PPIs) have been identified using a variety of technologies, including yeast two hybrid assays and immunoprecipitation-mass spectrometry (Luck et al. 2020; Yugandhar et al. 2019). Although these approaches are resource intensive, most human PPIs have been evaluated using large experimental efforts and cataloged in publicly available databases (Bajpai et al. 2020; Luck et al. 2020).

Because of the cost, a survey of all PPIs in various cell types is not feasible. Further, PPIs do not provide information on the cellular consequences of the relevant interactions. For example, proteins may have no physical interaction, even though their genes show strong interactions.

Genetic interactions have been evaluated in a wide variety of organisms, ranging from bacteria to Drosophila (Costanzo et al. 2019). The most thorough catalog is for the yeast *Saccharomyces cerevisiae* (Costanzo et al. 2016). This network has provided surprising new information on the connections between cellular pathways. Data in other organisms is far less complete. Assuming 20 000 protein coding genes in mammals, the number of potential interactions is ~ 2 × 10^8^. The addition of non-coding genes trebles the number of genes, bringing the number of possible interactions to ~ 1.8 × 10^9^.

Multiple opportunities for therapeutic intervention in cancer will emerge from genetic interaction maps (Mair et al. 2019). In particular, geneticists are beginning to use CRISPR-Cas9 genetic editing technology to study the viability of cancer cells with double loss-of-function mutations in *trans*. One recent study employed CRISPR interference (CRISPRi) to identify genetic interactions among 222 784 gene pairs in two cancer cell lines (Horlbeck et al. 2018). However, even this effort only evaluated ~ 0.1% of all possible coding gene pairs.

Overexpression alters cell physiology differently to loss-of-function, and is a complementary strategy to understanding cellular networks in both yeast and mammalian cells (Khan et al. 2020; Prelich 2012; Sopko et al. 2006). One recent approach to obtaining targeted increases in gene expression is CRISPR activation (CRISPRa), but this method causes constitutive and non-physiological overexpression (Gilbert et al. 2014; Kampmann 2018).

Radiation hybrid (RH) mapping has been widely used to construct genetic maps for the genome projects (Avner et al. 2001; Cox et al. 1990; Goss, Harris 1975; Hudson et al. 2001; Kwitek et al. 2004; McCarthy 1996; McCarthy et al. 2000; Olivier et al. 2001; Walter et al. 1994). In this technique, lethal doses of radiation are used to randomly fragment the genome of a human cell line (**Fig. 1A**) (Goss, Harris 1975). RH clones are then created by transferring a sample of the DNA fragments to living hamster cells using cell fusion. Linked markers are likely to reside on the same fragment, and hence co-inherited in the RH clones. Because of the small size of the DNA fragments, genotyping a panel of RH clones allows high resolution mapping.

**Figure 1.**
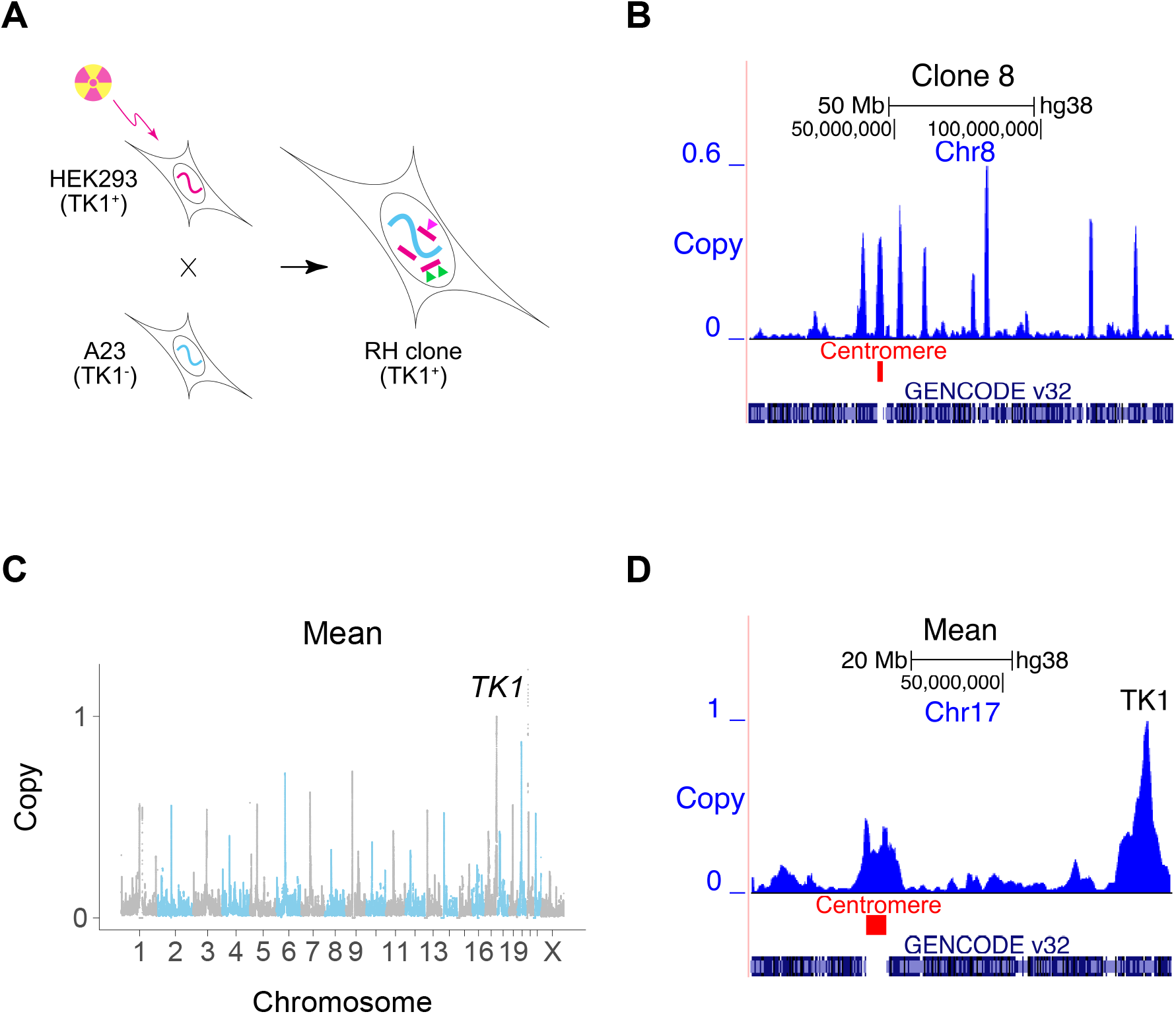
Creating RH interaction networks. (**A**) RH clones are made by lethally irradiating human cells (HEK293). The human cells are fused to living hamster cells (A23) to rescue the DNA fragments, and RH clones selected using *TK1*. Conventional RH mapping exploits the fact that nearby makers tend to be found on the same DNA fragment and thus co-inherited (green triangles). RH interaction networks seek co-inheritance of distant markers (green and pink triangles). (**B**) Human DNA copy number, clone 8, Chromosome 8. One of the human fragments encompasses the centromere. (**C**) Mean retention across 15 clones, showing increased retention at centromeres and retention of 1 at *TK1*. (**D**) Mean retention across clones, Chromosome 17.

We previously used publicly available RH data to construct a genetic interaction map for mammalian cells (Lin et al. 2010; Lin, Smith 2015). We reasoned that significant co-inheritance of marker pairs separated from each other in the human genome would disclose genetic interactions promoting cell survival. Rather than loss-of-function, this interaction network depends on extra gene copies. Moreover, the genes are expressed using their natural promoters instead of constitutively.

We increased the statistical power of the RH network by combining PCR genotyping datasets from six RH panels: three human, one mouse, one rat and one dog. The network consisted of 18 324 genes linked by 7 248 479 interactions. The overwhelming majority of interactions consisted of higher than expected co-retention of the gene pairs.

Despite its statistical power, the quality of the network suffered because of the need to combine datasets from different species. In addition, the mapping resolution was limited by the legacy PCR genotyping.

In this study, we demonstrate the cost-effectiveness of the RH approach in creating genetic interaction maps for the mammalian genome. We used low pass DNA sequencing to genotype emergent human RH clones, decreasing labor and cell culture costs, while also improving mapping resolution. Expanding this strategy will render construction of whole genome interaction networks feasible.

## Results

### Human DNA retention in the RH clones

We exploited the sensitivity of modern DNA technologies to analyze RH clones upon appearance, avoiding the customary need for additional growth. Our strategy cuts costs and saves time. Fifteen independent RH clones were evaluated using low pass sequencing at a depth of 0.31 ± 0.03 times the human genome (14.5 ± 1.4 M single reads of 65 bp). Reads were aligned to the human genome and those that also aligned to hamster were discarded. Only human-specific reads remained for subsequent analyses using 1 Mb windows and 10 kb steps (Khan et al. 2020).

The human DNA fragment length was 2.3 ± 0.1 Mb and the retention frequency was 0.25 ± 0.08 (**Figs. 1B–D**, **S1A–D**). Due to the small number of clones, 3 % of the human genome had zero retention. Clone 2 had the highest retention (0.97) and harbored a nearly complete copy of the human genome, plus additional fragments (**Fig. S2**). The human genome was represented with 3.8-fold ±1.3 redundancy in the 15 clones.

Centromeres showed increased retention, since they stabilize the donated human DNA fragments (Khan et al. 2020; Wang et al. 2011) (**Figs. 1B–D**, **S1E**). Fragments containing centromeres were also significantly longer than other fragments, due to the large size of these chromosomal elements (**Fig. S1F**). The human *TK1* gene was used as the marker to select hybrids, and its copy number was therefore 1.

### Mapping accuracy

Based on the average fragment length, retention frequency and panel size, the expected mapping resolution was 0.3 ± 1.2 Mb. We further estimated mapping resolution by evaluating the distance at which peak −log_10_ *P* values for *cis* linked 1 Mb windows decreased by one (−1 log_10_ *P* values) (**Fig. S3**). The −1 log_10_ *P* value was 1.0 Mb using −log_10_ *P* values plotted against distance and 2.2 Mb using distance against −log_10_ *P*.

As a surrogate measure of mapping accuracy, we also evaluated the distance between the retention peak of *TK1* and its known location (Khan et al. 2020). We did the same for the centromeres. The mapping resolution for *TK1* was 33.3 ± 59.5 kb and for the centromeres, −0.5 ± 2.5 Mb (**Fig. S4**), neither of which were significantly different from the expected value of zero (*P* = 0.06 and 0.80, respectively).

Although there were substantial differences between the mapping accuracy estimates for this small RH panel, the resolution is probably of the order of ≲ 1 Mb.

### Human genetic interactions

We identified human genetic interactions by seeking co-inheritance of DNA fragments > 2.4 Mb apart, using Fisher’s exact test to assess significance (false discovery rate, FDR < 0.05) (**Fig. 2**). Interacting genes were identified as genes closest to a −log_10_ *P* peak consisting of a single window pair, with less significant neighboring window pairs.

**Figure 2.**
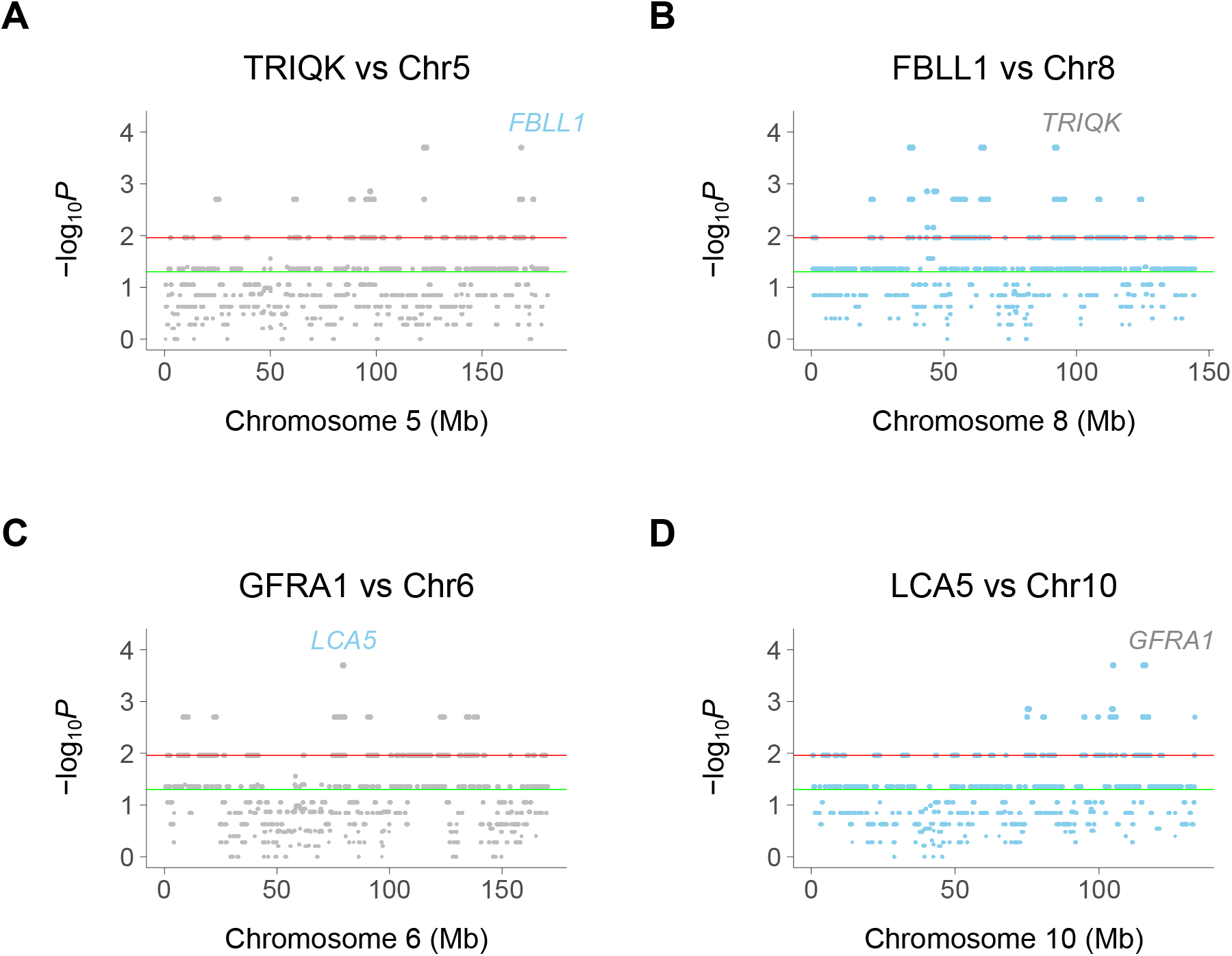
Chromosome plots of human genetic interactions. (**A**) Co-inheritance of peak window at *TRIQK* and windows on Chromosome 5, with *FBLL1* exceeding significance thresholds. (**B**) Co-inheritance of *FBLL1* and windows on Chromosome 8, with *TRIQK* exceeding significance thresholds. (**C**) Co-inheritance of *GFRA1* and windows on Chromosome 6, with *LCA5* exceeding significance thresholds. (**D**) Co-inheritance of *LCA5* and windows on Chromosome 10, with *GFRA1* exceeding significance thresholds. Green horizontal line, *P* = 0.05, Red horizontal line, FDR = 0.05.

Because of the selection procedure for interaction peaks, which ignores “plateaus” of −log_10_ *P* values, the realized mapping resolution of the network may be better than our empirical estimates. In fact, the resolution may be of the order of the 10 kb steps, close to the estimate using *TK1* retention.

We restricted our analysis to coding region genes to facilitate comparison with the legacy RH-PCR network as well as PPI networks. A total of 2359 interactions connecting 225 genes were found in the RH-Seq network, with a mean degree of 21.0 ± 1.4 (**Figs. 3A,B** and **S5**).

**Figure 3.**
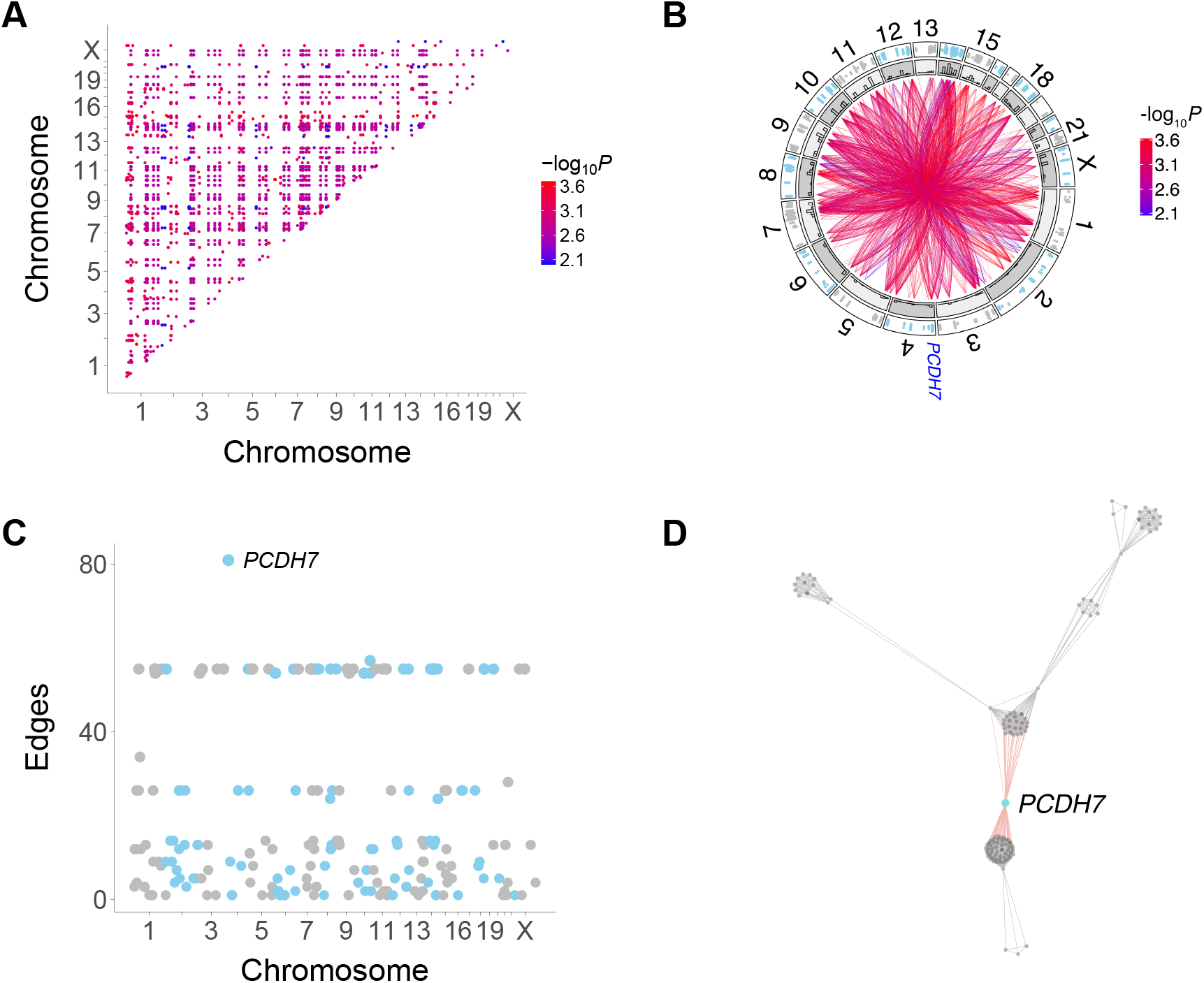
Human genetic interactions. (**A**) Location and significance of interactions. (**B**) Circos plot of interactions. Outer track, location of interacting genes; inner track, histograms of interaction numbers normalized for each chromosome. Location of PCDH7 on Chromosome 4 shown. (**C**) Number of interactions for each gene. *PCDH7* has the most. (**D**) Subnetwork featuring *PCDH7*.

All interactions in the RH-Seq network were “attractive”, in which gene co-retention occurred more often than expected by chance. Gene pairs that interact as extra copies thus promote cell growth. The interactions in our original RH-PCR network were also overwhelmingly attractive (Lin et al. 2010). This finding contrasts with loss-of-function alleles in yeast and cancer cells, where some allele combinations promoted, while others inhibited, growth (Costanzo et al. 2016; Costanzo et al. 2019; Horlbeck et al. 2018).

*PCDH7* had the largest number of interactions in the RH-Seq network, 81 (**Figs. 3B–D**). In addition, *PCDH7* regulated the expression of the most genes (614) in a mouse RH panel (Ahn et al. 2009; Park et al. 2008). Perhaps genes with many interactions also regulate the expression of many genes.

### Network overlaps

We found highly significant overlap between the RH-Seq interaction network and the legacy RH-PCR network (**Figs. 4A,B**) (maximum overlap, FDR *<* 2.2 × 10^*−*16^, odds ratio = 3.1, 97 common interactions, 32 expected). Considering the limited power of the RH-Seq dataset, the overlap is encouraging.

**Figure 4.**
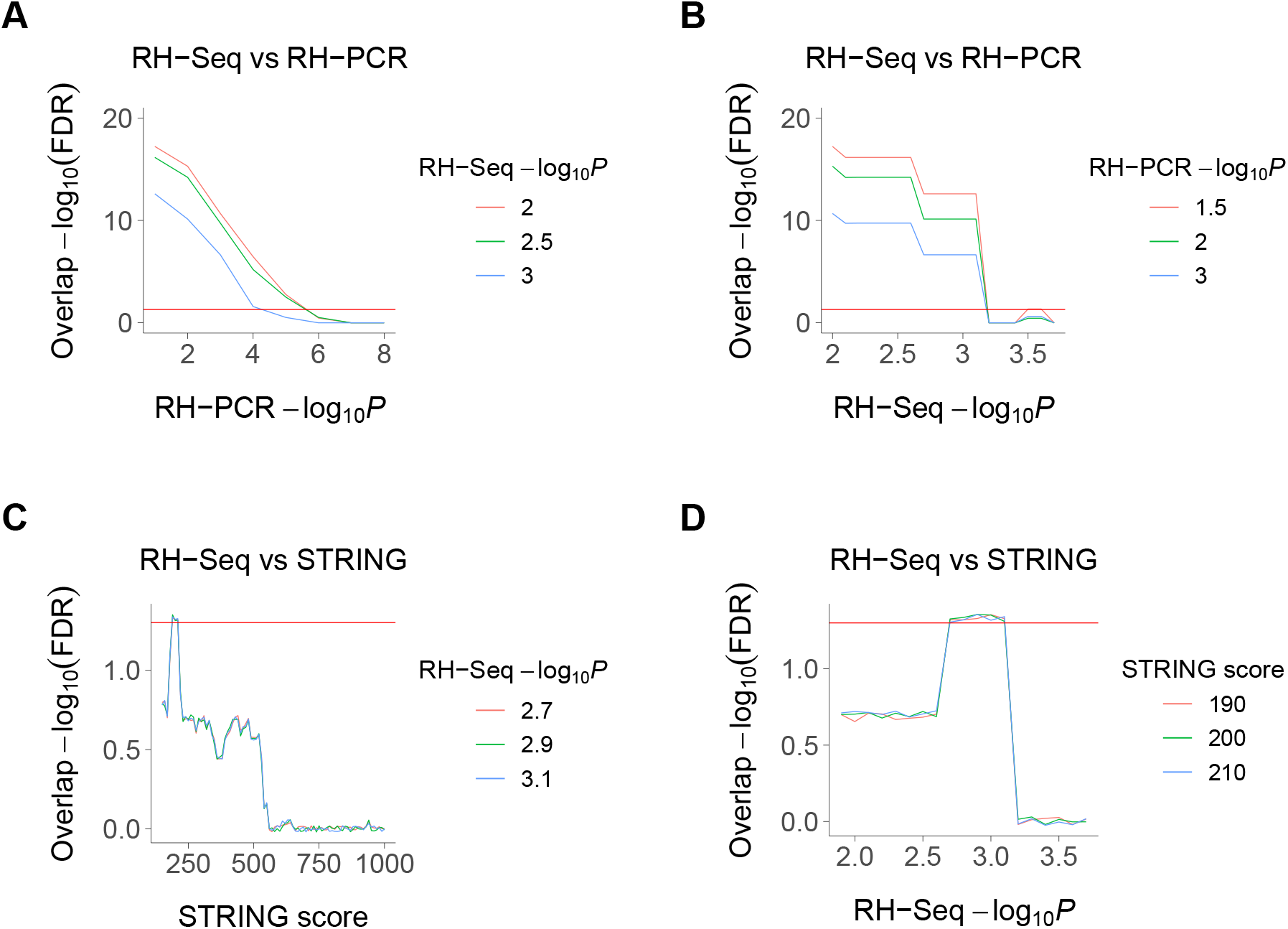
Overlaps of RH-Seq network. (**A**) Overlap of RH-Seq and RH-PCR networks, thresholded on RH-PCR −log_10_ *P* (abscissa). Overlap significance diminishes as threshold increases due to decreased numbers. (**B**) Overlap of RH-Seq and RH-PCR networks, thresholded on RH-Seq −log_10_ *P*. (**C**) Overlap of RH-Seq and STRING interactions, thresholded on STRING score. (**D**) Overlap of RH-Seq and STRING interactions, thresholded on RH-Seq − log_10_ *P*. Horizontal red lines, FDR = 0.05.

We also found less strong, but still significant, overlap between the RH-Seq interaction network and the STRING v11 PPI database (maximum overlap, FDR = 4.6 × 10^*−*2^, odds ratio = 2.3, 25 common interactions, 11 expected) (**Figs. 4C,D**) (Szklarczyk et al. 2019). We found no overlap with four other PPI datasets, BioGRID, HIPPIE, HINT and a yeast two-hybrid dataset, reflecting the modest size of the RH-Seq network (Alanis-Lobato, Schaefer 2020; Bajpai et al. 2020; Das, Yu 2012; Luck et al. 2020; Oughtred et al. 2021).

Nevertheless, the overlap of the RH-Seq and STRING networks suggests that gene products which interact to promote cell growth also tend to interact physically. A similar overlap was found between loss-of-function genetic interactions using CRISPRi in two cancer cell lines and the STRING database (Horlbeck et al. 2018). There was no overlap between the RH-Seq and the CRISPRi networks, but it is difficult to draw firm conclusions about the similarities and differences of these networks given that each are of limited size.

The original RH-PCR interaction network showed significant overlap with networks constructed using a gene-disease database and using GO (Gene Ontology Consortium 2021; Lin et al. 2010; Pletscher-Frankild et al. 2015). We found that the RH-Seq network also overlapped with these networks (**Fig. S6**). Interacting genes cause similar diseases and have similar functions.

### RH-Seq network and growth genes

We recently used selection of pooled RH cells to identify mammalian growth genes (Khan et al. 2020). Consistent with the idea that the RH-Seq interaction network is involved in control of cell proliferation, there was significant overlap between the RH-Seq network and the RH growth genes (**Figs. 5A,B**). In contrast, both the RH-Seq network genes and RH growth genes (Khan et al. 2020) showed significant non-overlap with growth genes identified in CRISPR loss-of-function screens, depending on the cell type (Hart et al. 2015; Wang et al. 2015) (**Figs. 5C–F**). These observations support the idea that over-expression alters cell physiology differently from loss-of-function, and suggests that the two approaches will give complementary interaction networks.

**Figure 5.**
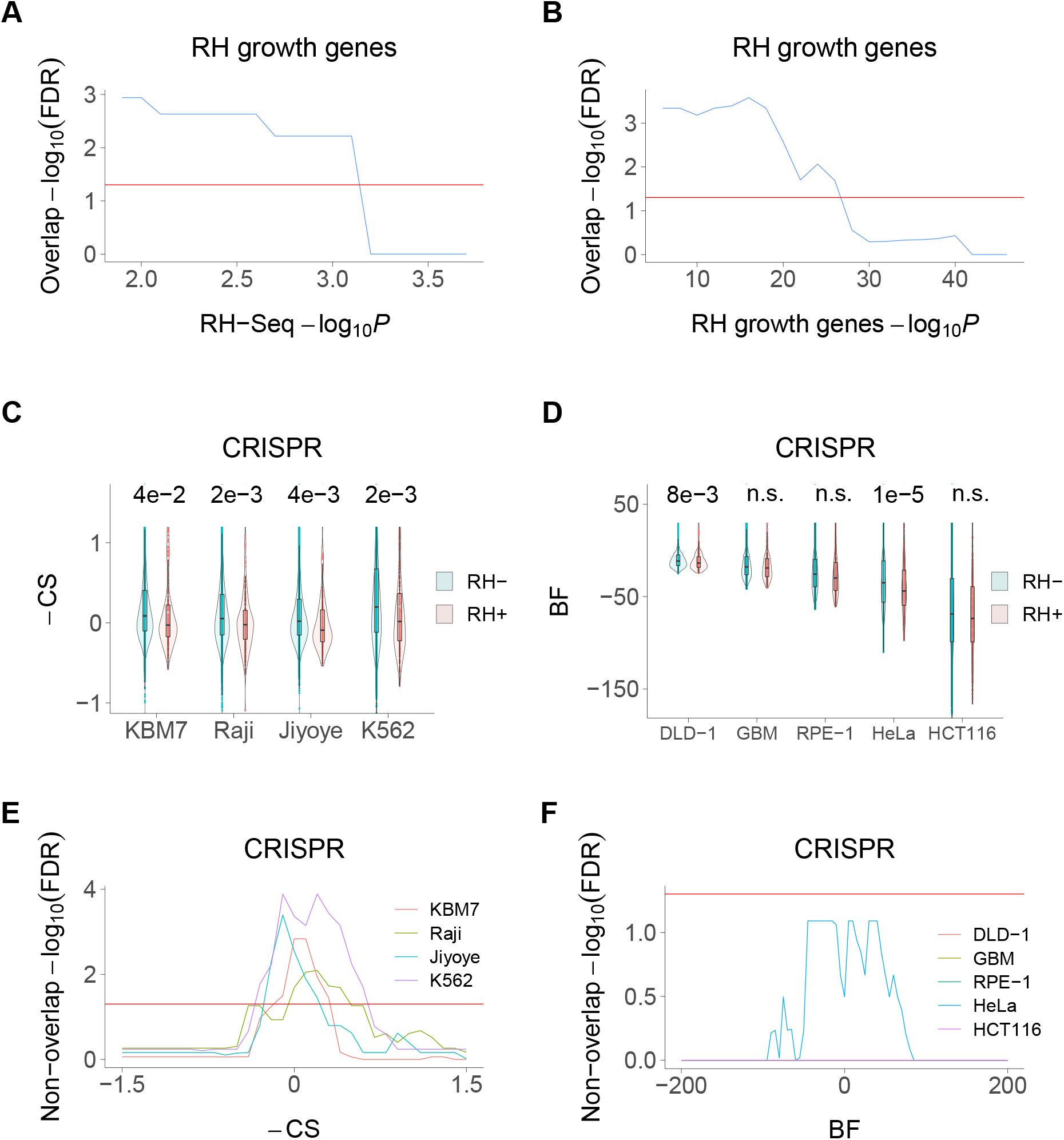
RH-Seq network and growth genes. (**A**) Overlap of RH-Seq network and RH growth genes, thresholded on RH-Seq network log10 *P*. Overlap significance diminishes as threshold increases due to decreased numbers. (**B**) Overlap of RH-Seq network and RH growth genes, thresholded on RH growth gene −log_10_ *P*. (**C**) RH-Seq interaction genes (RH+) show lower growth effects when inactivated using CRISPR than non-interaction genes (RH−). Higher –CS values (CRISPR score times −1) mean stronger growth effects of CRISPR null alleles (Wang et al. 2015). *P* values shown above comparisons. (**D**) RH-Seq interaction genes show lower growth effects when inactivated using CRISPR. Higher Bayes factors (BF) means stronger growth effect of CRISPR null alleles (Hart et al. 2015). (**E**) Significant non-overlap of RH-Seq network genes and CRISPR growth genes, thresholded on –CS score. (**F**) Non-overlap of RH-Seq network genes and CRISPR growth genes, thresholded on BF score. Horizontal red lines, FDR = 0.05.

### RH-Seq network and expression

The genes in the RH-Seq network showed decreased expression in a human RH panel (**Fig. S7A**), but not in the GTEx dataset (v8) of human tissue expression (**Fig. S7B**) (GTEx Consortium 2020; Wang et al. 2011). Similarly, the RH growth genes showed decreased expression in both the human RH panel and the GTEx dataset (Khan et al. 2020). Genes identified for their interactions and growth effects as a result of an extra copy are likely to be evolutionarily selected for decreased expression.

### Functional enrichment

The interacting genes in the RH-Seq network were highly enriched in cancer terms from a disease database (FDR < 0.05), with the top eight terms being cancer related (**Figs. S8A,B** and **S9A**) (Kuleshov et al. 2016; Pletscher-Frankild et al. 2015). This observation is consistent with a key role for the RH-Seq network in cell proliferation. Examples of cancer related genes in the network included *GAS6*, *MAP3K4*, *PRKD1*, *PTPRD* and *VEGFA*.

The RH-Seq interaction network was also significantly enriched in the catalog of human genome-wide association studies (GWAS) (Kuleshov et al. 2016) and in the NCBI Database of Genotypes and Phenotypes (dbGaP; https://www.ncbi.nlm.nih.gov/gap/) (FDR < 0.05) (**Figs. S8C** and **S9B,C**). The same was true for the RH growth genes (Khan et al. 2020). In contrast, growth genes from loss-of-function CRISPR screens showed no such enrichment. The effects of an extra gene copy in the RH cells may be closer to the mild effects of common disease variants than the more severe effects of a knockout.

Gene ontology analysis (GO) of the genes in the RH-Seq network revealed enrichment in a number of categories related to cell growth (*P* < 0.05, but FDR > 0.05), including cell proliferation, p38MAPK, growth factor, tyrosine kinase and replication fork (**Figs. S10** and **S11**) (Gene Ontology Consortium 2021).

### Evolutionary properties of RH-Seq network genes

The RH-Seq network genes displayed significantly decreased numbers of duplications (paralogs) (**Fig. S12A**). The evolutionary selection against duplication of the RH-Seq network genes is consistent with their effects on cell survival as an extra copy. Similarly, the RH growth genes also showed decreased numbers of paralogs (Khan et al. 2020).

The RH-Seq network genes exhibited increased evolutionary conservation, with decreased tolerance to loss-of-function (LOF) variants (**Fig. S12B**) and decreased mouse-human sequence divergence (**Fig. S12C**). However, there was no significant increase in the number of species with orthologs of the network genes (“phyletic retention”), another measure of evolutionary conservation (**Fig. S12D**). The RH growth genes also displayed increased evolutionary conservation (Khan et al. 2020). Both the RH-Seq network genes and the RH growth genes had increased gene lengths (**Figs. S12E–S12H**).

## Discussion

We created a genetic interaction network using 15 RH clones. Gene interactions were identified by seeking unlinked gene pairs that showed significant co-retention. We used low pass sequencing of nascent RH clones to save labor and consumable costs, while obtaining high quality genotyping.

Unlike approaches such as weighted correlation network analysis (WGCNA), which use similarity of gene expression to assign function (Langfelder, Horvath 2008), the endpoint of the RH approach is cell proliferation. The RH network therefore plays a causative, rather than correlative, role in cell viability.

The RH-Seq network showed highly significant overlap with the original RH-PCR network as well as less strong, but still significant, overlap with a PPI database. The large number of potential interactions means that significance can occur even with modest overlaps. Nevertheless, considering the small size of the RH-Seq dataset, the significant overlaps suggest that this strategy is a reproducible and scalable approach to genetic interaction networks.

The RH-Seq network genes displayed significant overlap with RH growth genes but non-overlap with growth genes identified using CRISPR loss-of-function screens. The extra gene copies used by the RH approach will provide complementary insights into cell physiology compared to loss-of-function networks.

The interacting RH-Seq genes were enriched in terms related to cancer in a gene-disease database, suggesting that the RH network strategy is relevant to cell proliferation and may provide new therapeutic insights into tumorigenesis.

Both the RH-Seq interaction genes and the RH growth genes showed significant enrichment in the GWAS and dbGaP databases, while the CRISPR growth genes did not (Khan et al. 2020). Most variants that contribute to complex traits are in non-coding regions and affect gene expression (Gallagher, Chen-Plotkin 2018). The significant enrichment of the RH-Seq network genes in the GWAS and dbGaP databases may reflect the milder physiological effects of an extra gene copy driven by its natural promoter, compared to the more severe effects caused by CRISPR loss-of-function alleles or overexpression using CRISPRa.

It has been suggested that cataloging genetic interactions will help illuminate the “missing heritability” in complex traits (Costanzo et al. 2019). The enrichment of the RH-Seq network genes in the GWAS and dbGaP catalogs suggests that the RH genetic interaction networks will be particularly relevant to understanding complex traits.

The cost of RH-Seq for mapping genetic interactions compares favorably to other approaches. Comprehensive coverage of the protein encoding interactome using CRISPR would require ~ 20 000 separate experiments. The RH-Seq approach offers the cost savings of low pass sequencing and lack of clone propagation. Further, each RH clone harbors ~ 0.25 times the human genome and therefore evaluates multiple genes in each cell. In contrast, CRISPR strategies evaluate only a single gene in each cell. The additional layer of multiplexing in RH-Seq lends efficiency to the construction of genetic interactions maps. In fact, the size of the RH-Seq network obtained using 15 clones (225 genes) is comparable to a recent large CRISPRi study, which examined genetic interactions for < 450 genes in cancer cells (Horlbeck et al. 2018).

The consumable cost per clone is ≲ $50 in RH-Seq, making the cost of analyzing 1000 clones feasible. The labor costs of data acquisition are also far lower than the other strategies. One individual could easily create and analyze 1000 clones. In addition to high statistical power, such a panel would have mapping resolution of 4 ± 18 kb, sufficient to confidently map individual genes (Lin et al. 2010).

The cost advantages of the RH-Seq strategy will allow it to be applied to wider areas. Analyzing nascent clones identifies the genetic interactions necessary for initial viability. Further culture of the clones will allow the evolution of genetic interactions to be evaluated over time. Expanding the cell fusion reaction to isogenic human cell lines will allow genetic networks to be mapped in the presence or absence of oncogenic mutations. Although these more ambitious experiments will increase costs, using 1000 RH clones would only be 5 % of the consumable cost of CRISPR approaches.

An additional advantage of the RH-Seq approach is that non-coding genes can be incorporated into the genetic network on an equal footing with coding genes (Khan et al. 2020). This extension will not increase the cost of the RH-Seq network, but would increase the cost of a CRISPR network nine-fold. In addition, the RH-Seq approach can evaluate haploinsufficient genes, which are difficult to assess using loss-of-function methods.

Low pass sequencing of nascent RH clones is a realistic and affordable approach to constructing comprehensive genetic interaction maps of the human genome. The strategy can be scaled in a cost-effective fashion to evaluate genetic networks that contribute to cancer and other complex disorders, providing new therapeutic insights.

## Methods

### Cells

We used human HEK293 (*TK1*^+^) and hamster A23 cells (*TK1*^−^), each previously validated by low pass sequencing (Khan et al. 2020). Cell fusion was performed as described (Khan et al. 2020). Before fusion, A23 cells were grown in the presence or absence of bromodeoxyuridine (BrdU, 0.03 mg ml^−1^). We irradiated the HEK293 cells using 100 (Gray) Gy, with an expected fragment length of 4 Mb, close to the observed. After fusion, cells were plated at a dilution of 1:10 in selective HAT medium (100 *μ*M hypoxanthine, *μ*M aminopterin, 16 *μ*M thymidine; Thermo Fisher Scientific^®^). RH clones (*TK1*^+^) were picked at 3 weeks.

### Sequencing

DNA was purified from 24 clones (20 BrdU and 4 non-BrdU) using the Illumina Nextera™ DNA Flex Library kit, following manufacturer’s instructions. Human DNA quantities as low as 100 ng can be sequenced using this kit. Illumina sequencing employed 65 bp single reads.

We obtained human-specific reads by aligning reads to the GRCh38/hg38 human genome assembly (hg38.fa) at high stringency, allowing only one mismatch (Khan et al. 2020). Reads that also aligned to the Chinese hamster (*Cricetulus griseus*) genome assembly (RAZU01) (Rupp et al. 2018) were then discarded, leaving human-specific reads. Alignments were quantitated using the number of human-specific reads per 1 Mb window with 10 kb steps (Khan et al. 2020).

A total of 16.0 ± 1.3 M reads were obtained for each of the 24 clones. Based on retained human fragments, a total of 9 clones were duplicates (7 were duplicates of clone 1, and 2 were duplicates of clone 9), presumably due to cell dispersion before clone picking. The 15 independent clones were used to construct the interaction network. In the independent RH clones, there were 589 ± 135 reads in 1 Mb windows containing human fragments and 18 ± 3 in those without.

### Evaluating mapping accuracy

To measure mapping resolution, we used the significance of *cis* linkage for windows separated by < 20 Mb. Linear models were employed to estimate initial slopes relating the −log_10_ *P* values and the distances between the windows and vice versa. The slopes were then employed to quantitate −1 log_10_ *P* values.

We also used the human DNA retention profiles averaged across the 15 clones to evaluate mapping accuracy. The midpoints of retention peaks for *TK1* or the centromeres were taken as the estimated location of each element (Khan et al. 2020). The distance between the estimated and actual midpoint positions was the mapping resolution. Standard errors of the mean were assessed by bootstrapping.

### Genetic interactions

Relative copy numbers in the 1 Mb windows were calculated by normalizing human-specific sequence reads to those at *TK1* in each clone. *TK1* has a retention frequency of one, since it is the marker used to select RH clones. A window was deemed to harbor a human DNA fragment if log2(relative copy number + 1) > 0.2, corresponding to the upper 28th percentile of values and a relative copy number of 0.15.

Fisher’s exact test was used to identify pairs of windows separated by *>* 2.4 Mb (upper 49^th^ percentile of fragment lengths) and whose co-inheritance were significantly different from the null (FDR < 0.05). Significant window pairs were designated as representing a genetic interaction if the log10 *P* peak consisted of a single window pair, with adjacent window pairs showing decreased −log_10_ *P* values. The genes corresponding to the interaction peak were taken as the nearest protein coding genes using GENCODE v31 (Frankish et al. 2021).

### Networks

The gene-disease network was constructed using a literature based database (Pletscher-Frankild et al. 2015). Genes sharing the same disease were linked together, yielding a network with 16 254 genes and 23 983 208 interactions. The GO network was constructed by linking genes sharing the same GO categories containing ≥ 70 and≤ 1000 members (Lin et al. 2010). The network consisted of 16 902 genes and 9 581 750 interactions.

### Public data

GO analyses used the 2021-01-01 release and the PANTHER Overrepresentation Test, as well as Enrichr (Gene Ontology Consortium 2021; Kuleshov et al. 2016). The duplicated genes database (DGD) together with Ensembl release 71 was used to identify paralogs (Ouedraogo et al. 2012).

Intolerance of human genes to predicted loss-of-function (pLoF) variants was evaluated using the observed/expected ratio from the Genome Aggregation Database (gnomAD) release 2.1.1. (Karczewski et al. 2020). We assessed the evolutionary divergence of mouse-human homologs using the ratio of nonsynonymous to synonymous substitutions (dN/dS) in Ensembl release 97 (Cunningham et al. 2019). The number of species with gene orthologs was evaluated using Homologene release 68 (Sayers et al. 2019). Unless otherwise noted, tests of significance used Welch’s Two Sample t-test.

## Data access

The sequencing data generated in this study have been submitted to the NCBI BioProject database (https://www.ncbi.nlm.nih.gov/bioproject/) under accession number PRJNA714684. Other data, tables and computer scripts are available from figshare (https://figshare.com/; http://dx.doi.org/10.6084/m9.figshare.14481258).

## Competing interest statement

The authors declare no competing interests.

## Acknowledgments

We acknowledge the UCLA Semel Institute Neurosciences Genomics Core for sequencing. This work used computational and storage services associated with the Hoffman2 Shared Cluster provided by the UCLA Institute for Digital Research and Education Research Technology Group.

## Author contributions

AHK: Acquisition of data, analysis and interpretation of data, drafting or revising the article. DJS: Conception and design, analysis and interpretation of data, drafting or revising the article.

---

**Supplemental Figure S1.**
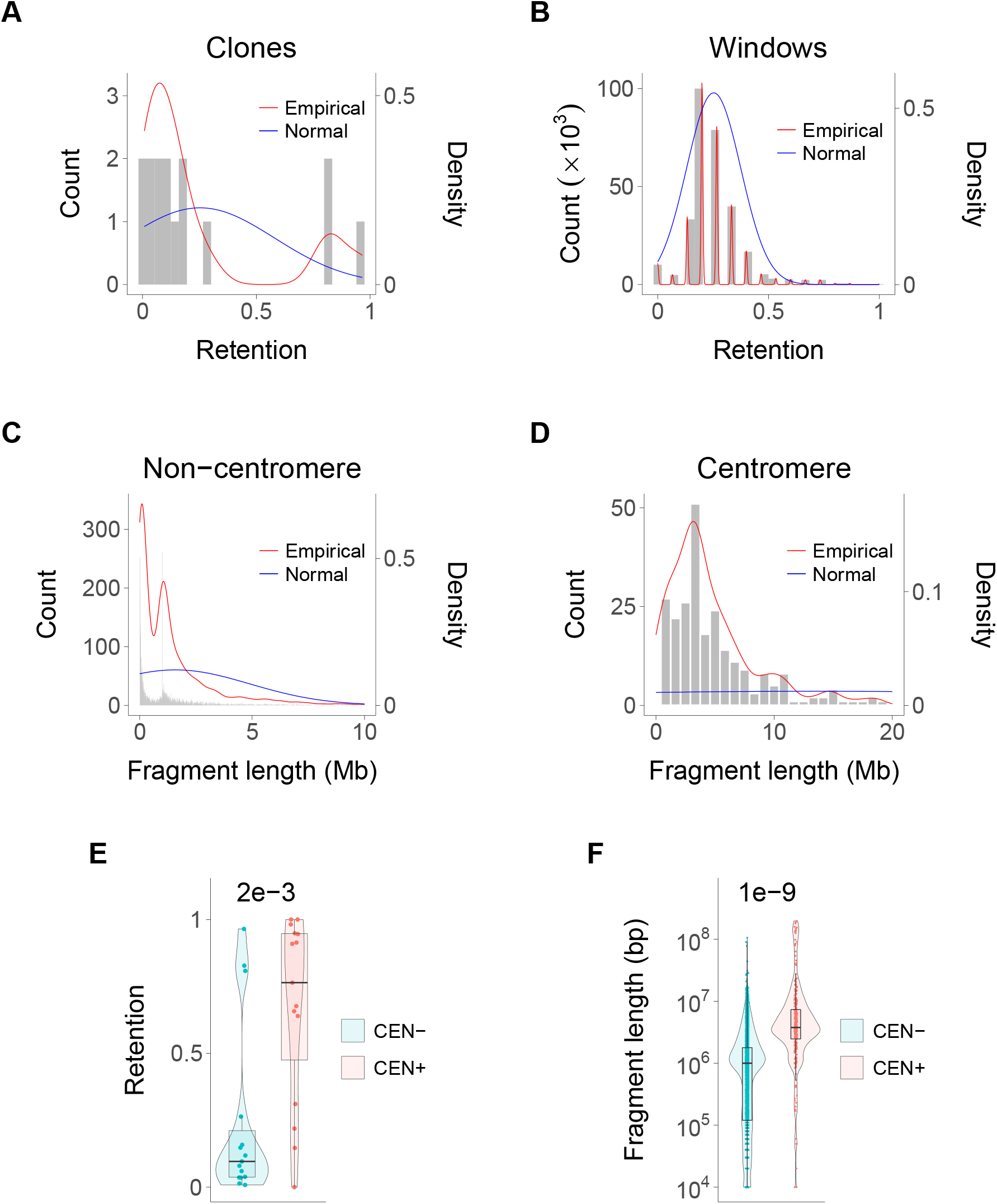
Human DNA in the RH clones. (**A**) Retention for each clone. (**B**) Retention for each 1 Mb window. (**C**) Non-centromeric fragment lengths. (**D**) Centromeric fragment lengths. (**E**) Retention of centromeric DNA (CEN+) is significantly higher than non-centromeric DNA (CEN−) (0.67 ± 0.09 vs 0.24 ± 0.09, *P* = 1.6 × 10^*−*3^). (**F**) Centromeric fragment lengths are significantly longer than non-centromeric (13.5 ± 1.9 Mb vs 1.6 ± 0.05 Mb, *P* = 1.4 × 10^*−*9^).

**Supplemental Figure S2.**
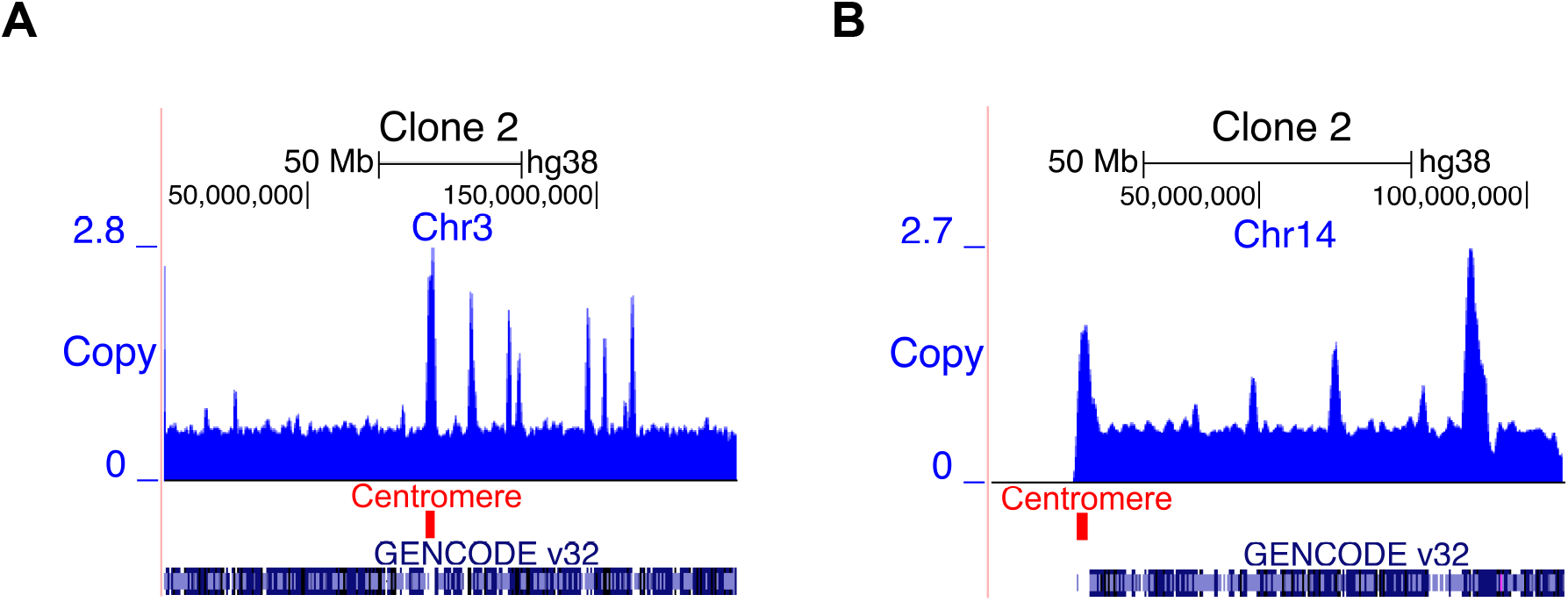
Human DNA retention in clone 2. (**A**) Human DNA copy number, clone 2, Chromosome 3. A complete human Chromosome 3 is present, in addition to smaller fragments. (**B**) Clone 2, Chromosome 14. A complete human Chromosome 14 is present, in addition to smaller fragments.

**Supplemental Figure S3.**
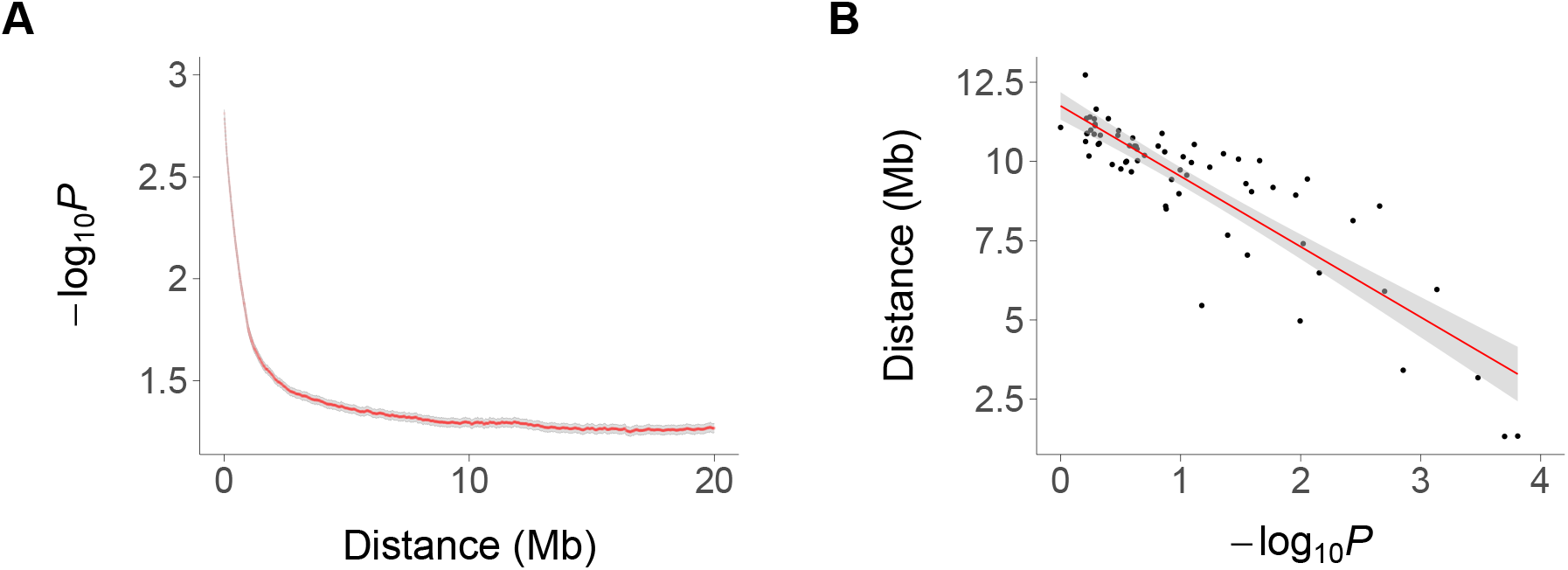
*Cis* linkage disequilibrium for RH-Seq clones. (**A**) Significance values for *cis* linked windows (− log_10_ *P*) plotted against distance. (**B**) Distance of *cis* linked windows plotted against − log_10_ *P*. Gray, 95% confidence intervals.

**Supplemental Figure S4.**
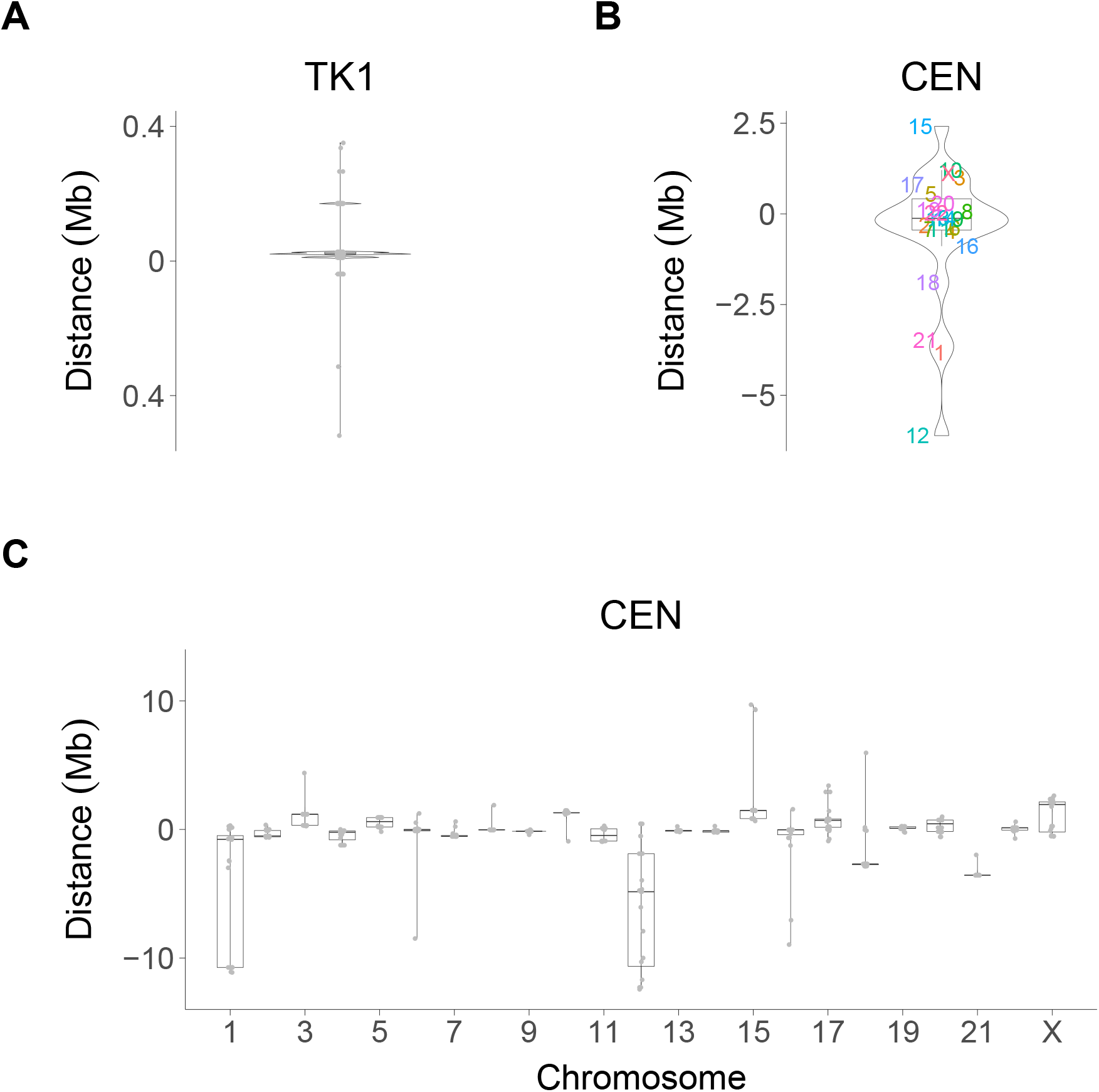
Mapping accuracy. (**A**) TK1. (**B**) Centromeres averaged across clones. Colored labels indicate chromosomes. (**C**) Centromeres vs chromosome. Bootstrapping used to evaluate variance.

**Supplemental Figure S5.**
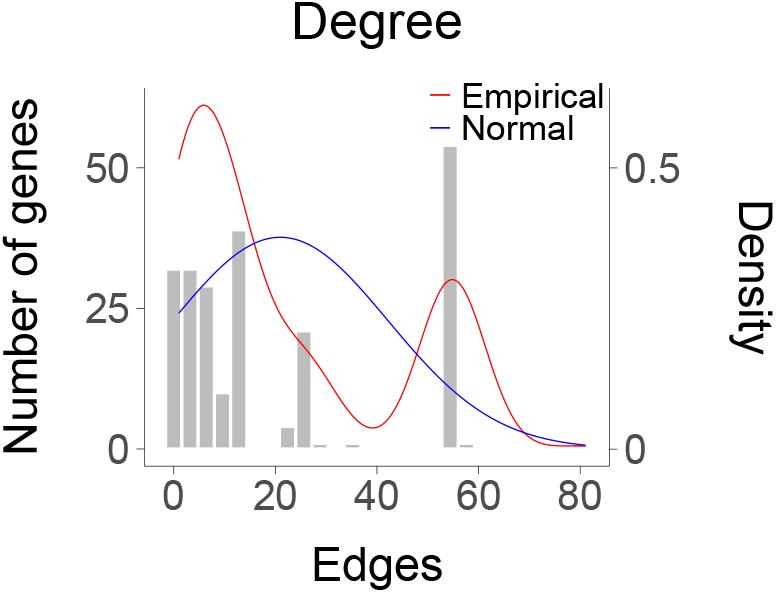
Degree of RH-Seq network.

**Supplemental Figure S6.**
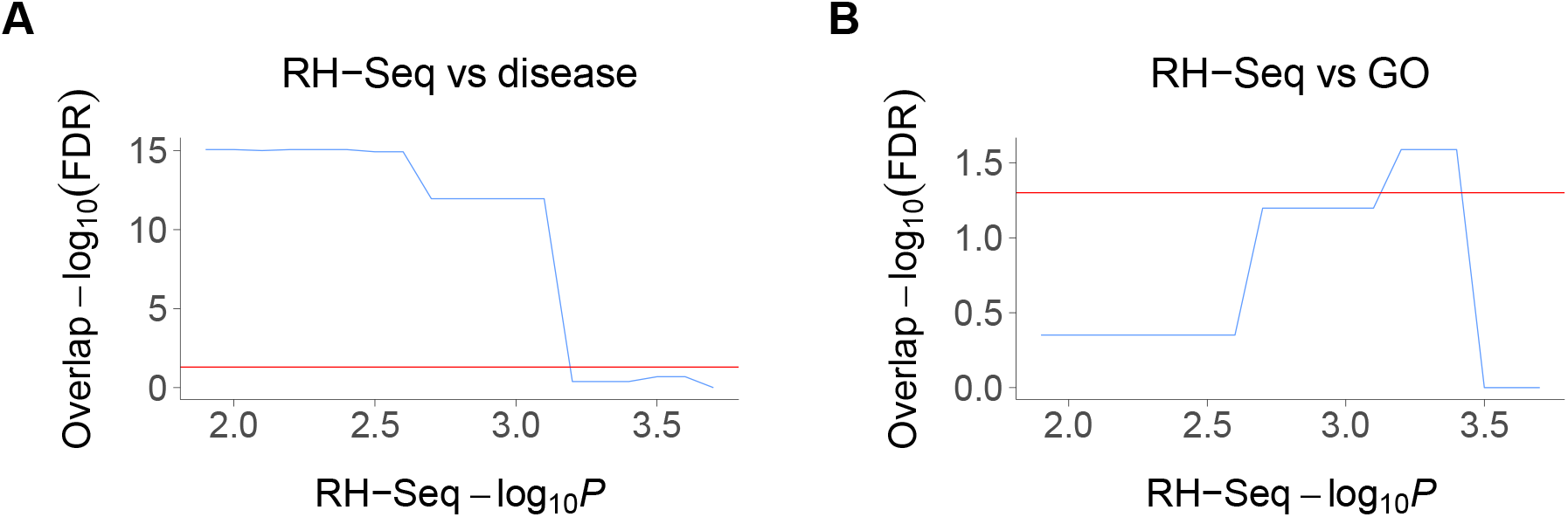
Overlaps between the RH-Seq network and gene-disease and GO networks. (**A**) Significant overlap of RH-Seq network and gene-disease network, thresholded on RH-Seq −log_10_ *P*. (**B**) RH-Seq network and GO network. Horizontal red lines, FDR = 0.05.

**Supplemental Figure S7.**
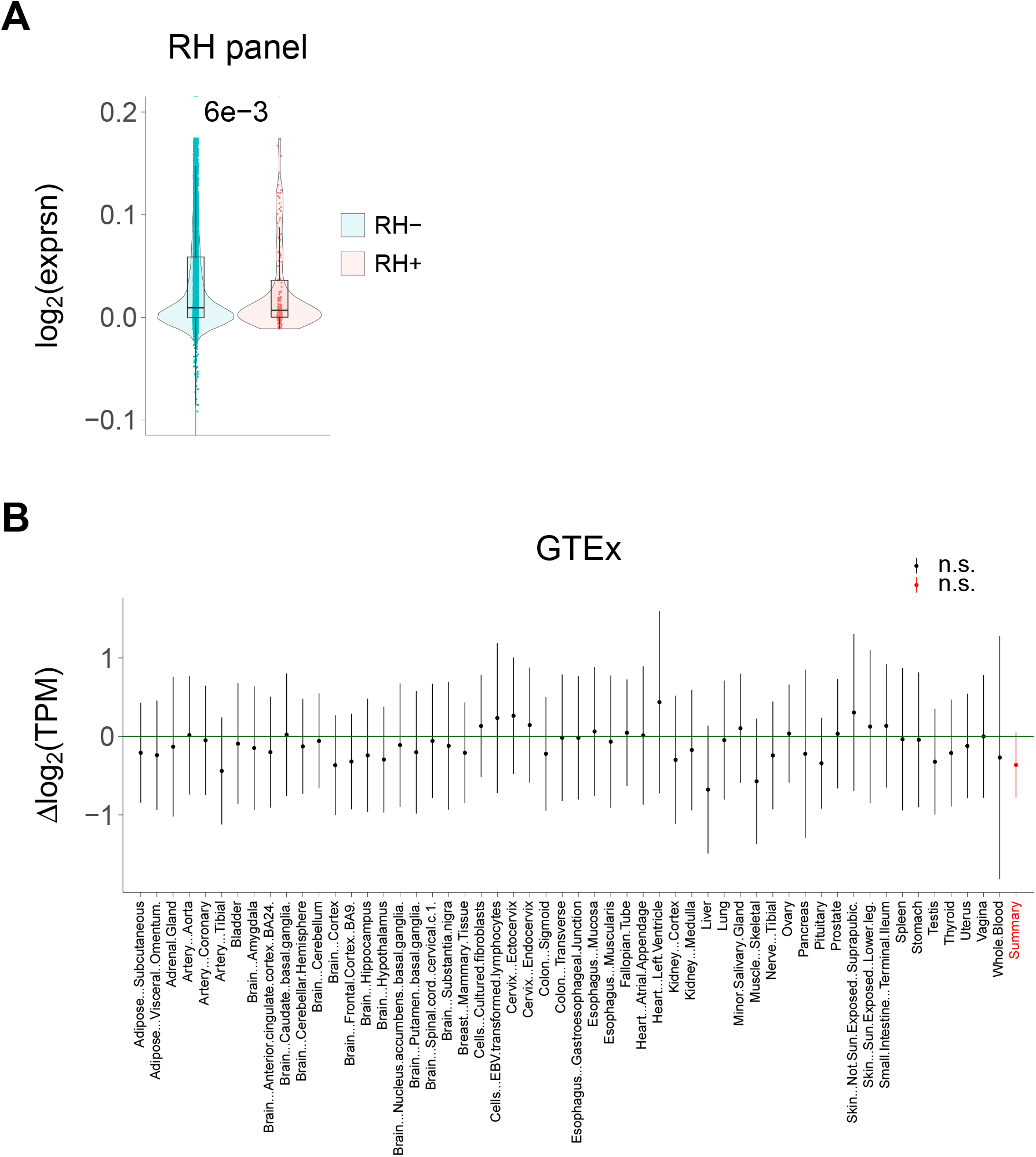
RH-Seq network and gene expression. (**A**) Genes in the RH-Seq network (RH+) show significantly lower expression than genes outside the network (RH−) using microarray data from a human RH panel (P = 5.7 × 10^−3^). (**B**) No significant differences in expression for RH-Seq network genes compared to non-network genes in various tissues using GTEx. Δlog_2_(TPM) < 0 means decreased expression for genes in RH-Seq network compared to non-network genes. Transcripts per million (TPM) thresholded at ≥ 5.

**Supplemental Figure S8.**
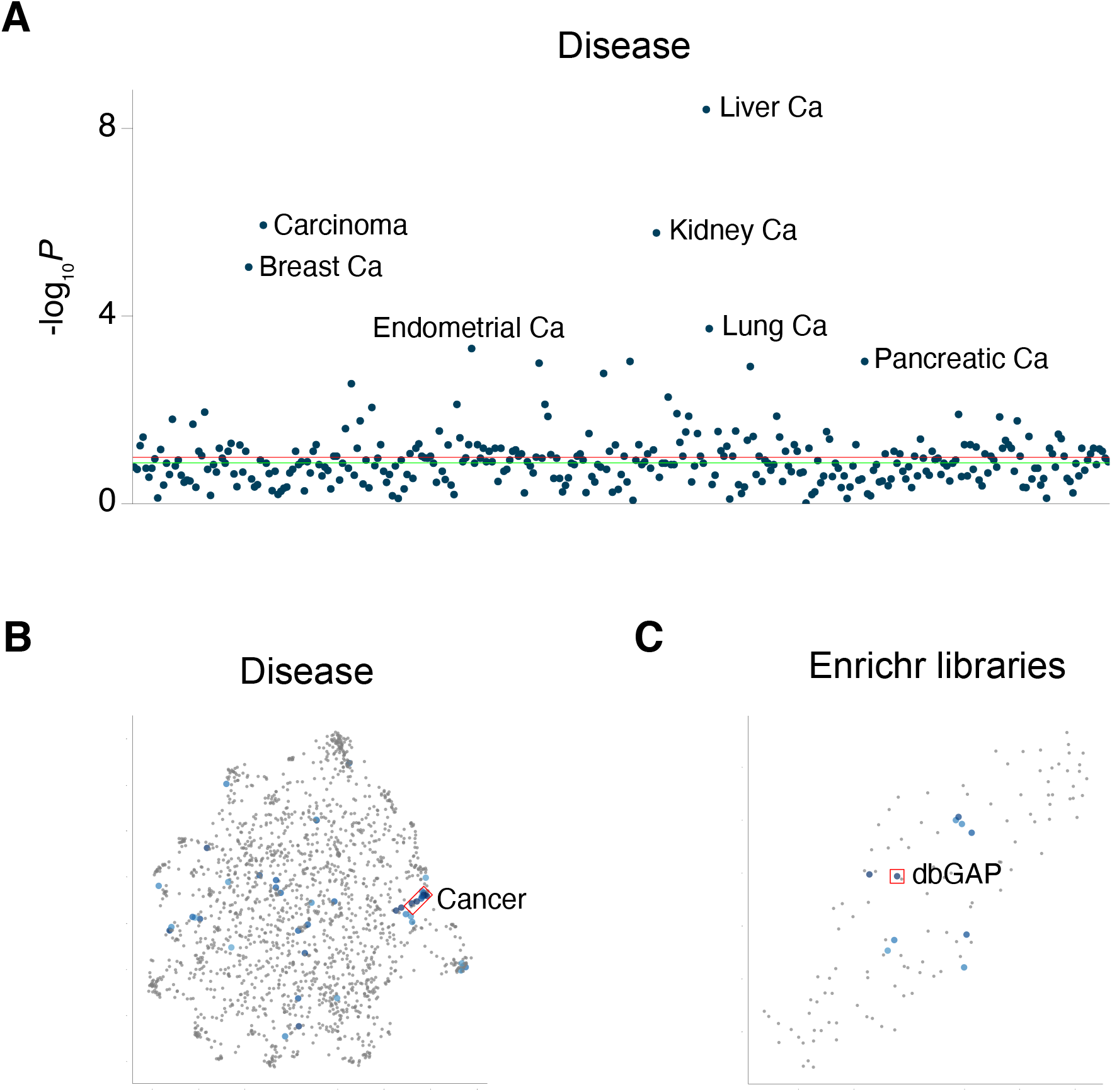
Enrichr analysis of genes in RH-Seq network. (**A**) Manhattan plot of human disease categories shows prominence of cancer. Green horizontal line, *P* = 0.05, red horizontal line, FDR = 0.05. (**B**) Scatterplot of human disease categories, showing significant over-representation of cancer. Related categories cluster together. Coordinates, arbitrary units. (**C**) Enrichr libraries with increased representation of human interaction genes, showing significant over-representation in dbGaP.

**Supplemental Figure S9.**
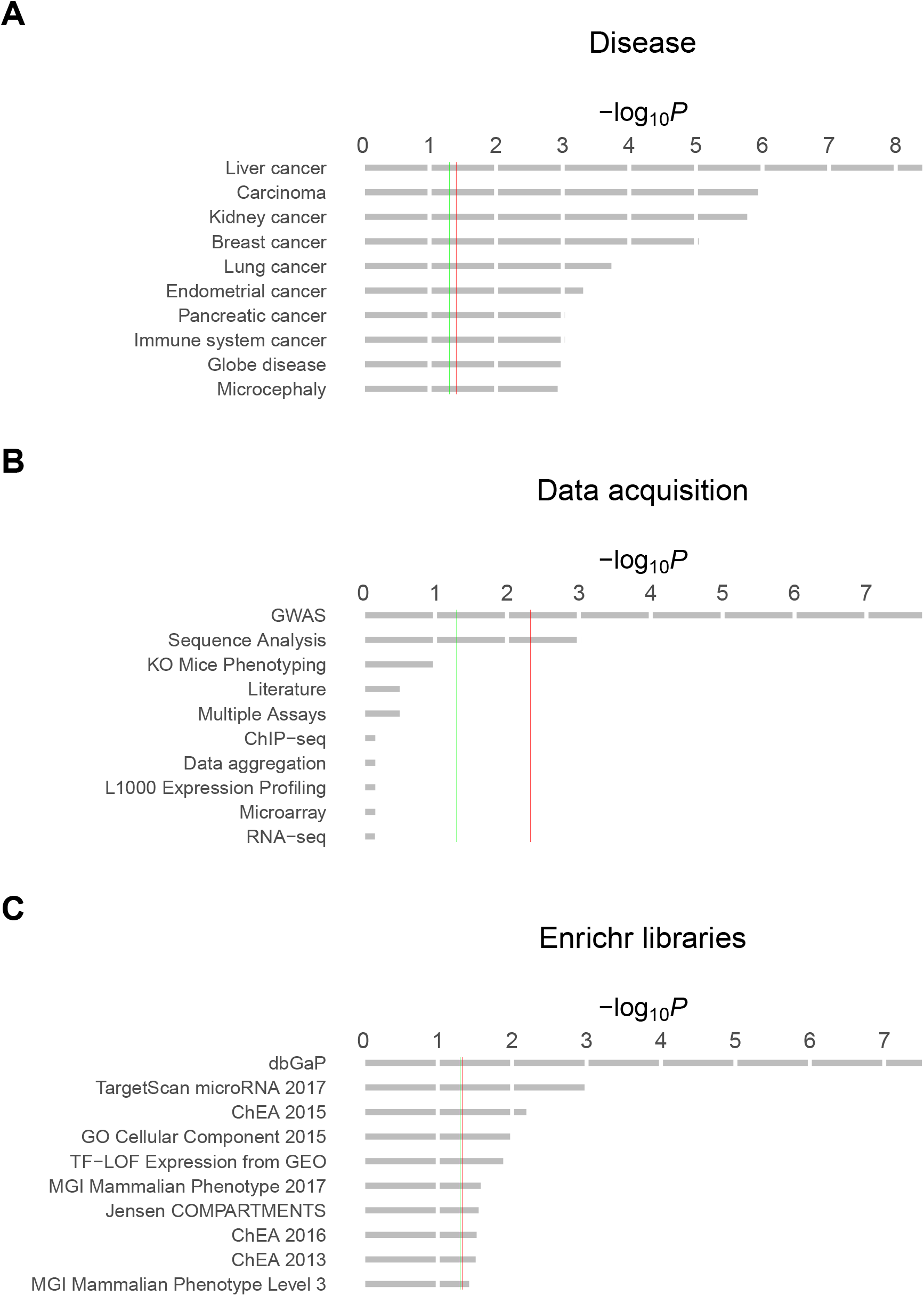
Enrichr analysis of genes in RH-Seq network, showing −log_10_ *P* values. (**A**) Disease categories, revealing prominence of cancer. (**B**) Most popular data acquisition method, with GWAS most significant. (**C**) Enrichr libraries, with dbGAP most significant. Green vertical lines, *P* = 0.05, red vertical lines, FDR = 0.05.

**Supplemental Figure S10.**
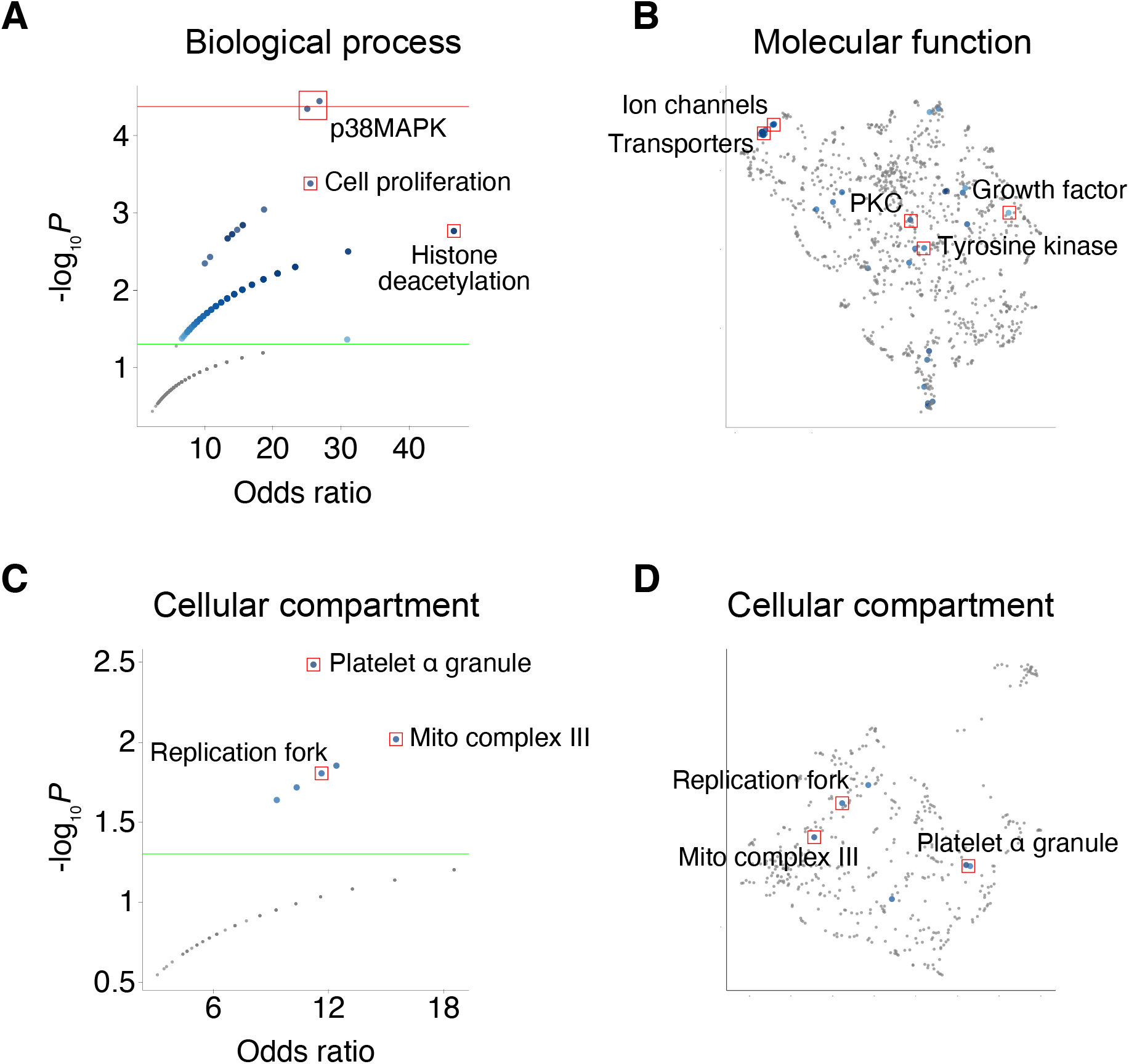
GO analysis of human genetic interactions using Enrichr. (**A**) Volcano plot of GO biological process. (**B**) Scatterplot of GO molecular function. Related categories cluster together. Coordinates, arbitrary units. (**C**) Volcano plot of GO cellular compartment. (**D**) Scatterplot of GO cellular compartment. Categories related to cell proliferation are enriched. Green horizontal lines, *P* = 0.05, red horizontal line, FDR = 0.05.

**Supplemental Figure S11.**
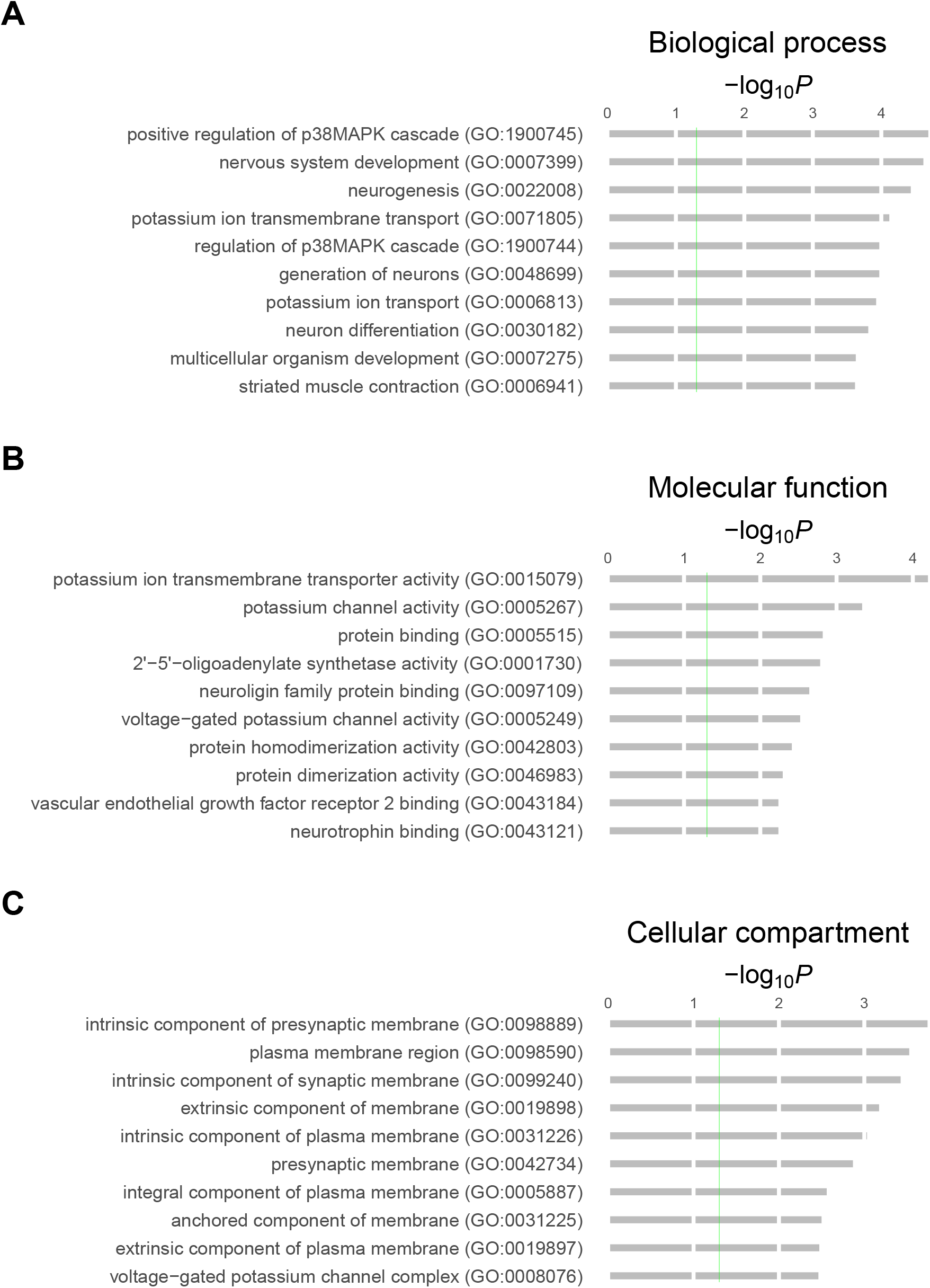
GO analysis of human genetic interactions using PANTHER Overrepresentation Test. (**A**) Biological process. (**B**) Molecular function. (**C**) Cellular compartment. Vertical green lines, *P* = 0.05.

**Supplemental Figure S12.**
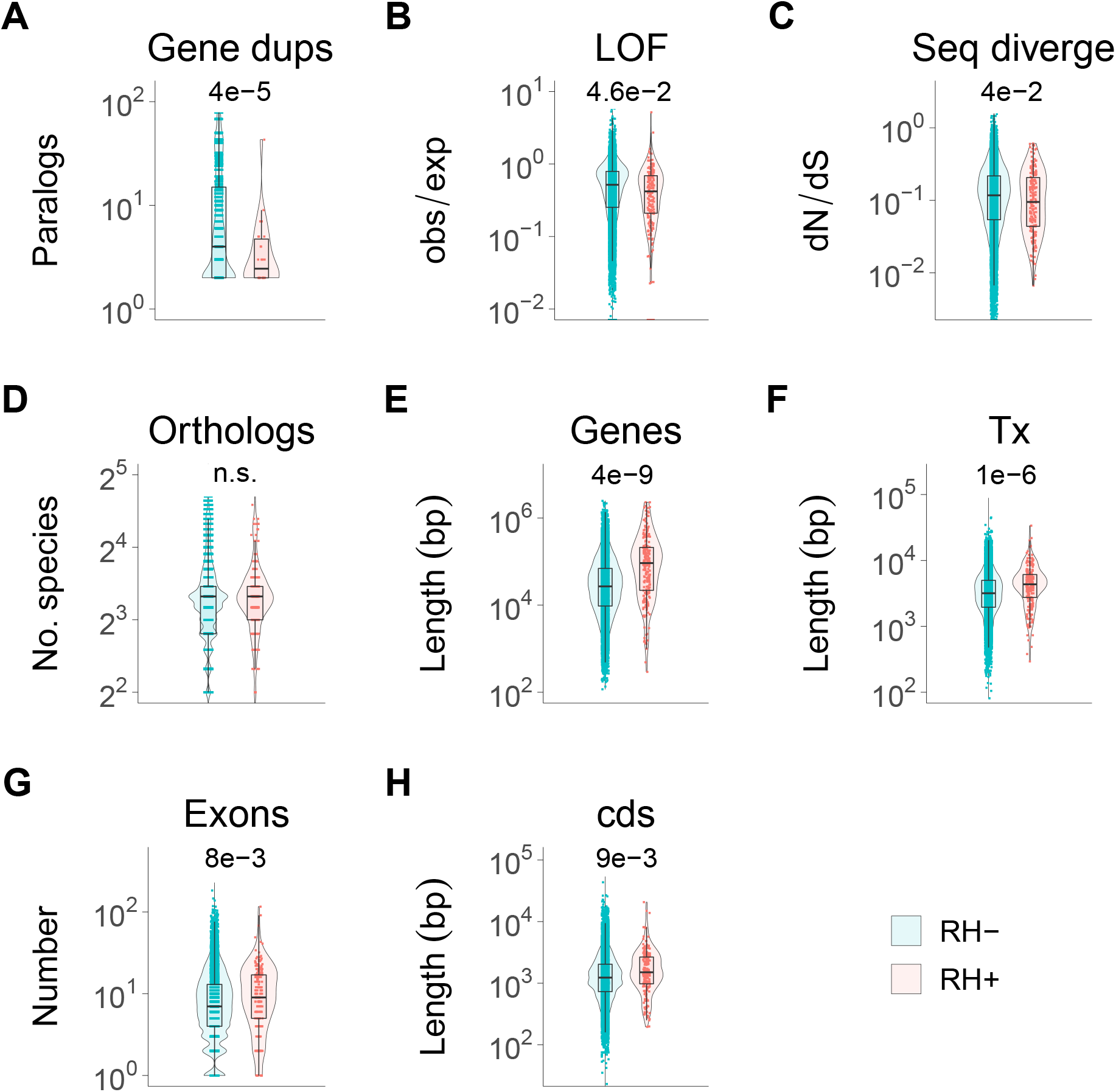
Evolutionary properties of RH-Seq network genes. (**A**) RH-Seq network genes show decreased gene duplications. (**B**) Decreased tolerance to loss-of-function (LOF) variants. (**C**) Decreased mouse-human sequence divergence. (**D**) No significant difference in the number of orthologs. (**E**) RH-Seq network genes show increased gene lengths. (**F**) Increased transcript lengths. (**G**) Increased exon numbers. (**H**) Increased coding sequence (cds) lengths. RH+, genes in RH-Seq network; RH−, genes outside network. *P* values shown above plots.

## Notes

https://www.ncbi.nlm.nih.gov/bioproject/?term=PRJNA714684

http://dx.doi.org/10.6084/m9.figshare.14481258

### Competing Interest Statement

The authors have declared no competing interest.

